# Cannabinoids modulate food preference and consumption in *Drosophila melanogaster*

**DOI:** 10.1101/2020.12.06.413914

**Authors:** Jianzheng He, Alice Mei Xien Tan, Si Yun Ng, Fengwei Yu

## Abstract

Cannabinoids have an important role in regulating feeding behaviors via cannabinoid receptors in mammals. Cannabinoids also exhibit potential therapeutic functions in *Drosophila melanogaster*, or fruit fly that lacks cannabinoid receptors. However, it remains unclear whether cannabinoids affect food consumption and metabolism in a cannabinoid receptors-independent manner in flies. In this study, we systematically investigated pharmacological functions of various cannabinoids in modulating food preference and consumption in flies. We show that flies display preferences for consuming cannabinoids, independent of their sensory functions. Interestingly, phyto- and endo- cannabinoids exhibit an inhibitory effect on food intake. Unexpectedly, the non-selective CB1 receptor antagonist AM251 attenuates the suppression of food intake by endocannabinoids. Moreover, the endocannabinoid anandamide (AEA) and its metabolite inhibit food intake and promote resistance to starvation, possibly through reduced lipid metabolism. Thus, this study has provided insights into a pharmacological role of cannabinoids in feeding behaviors using an adult *Drosophila* model.

## Introduction

Eating disorders including binge eating, and anorexia and bulimia nervosa, and obesity are characterized by abnormal eating behaviors and dysregulation of food intake. These have immense long-term consequences on physical health that could manifest to cardiovascular diseases, and on mental health that leads to both somatic and psychosocial complications ^1^. With the increasing burden on the society and healthcare system, research in the past decade has focused its efforts on understanding the mechanisms underlying the regulation of food intake and control of body weight, and ultimately developing therapeutic drug treatments to curb these issues.

*Cannabis sativa*, also known as marijuana, a herbaceous plant belonging to the family Cannabaceae, has been known for its medicinal and recreational purposes for over 5000 years ^2,3^. Recently, it has gained recognition as a valuable source of unique compounds with various potential therapeutic applications ^4-7^. It has since been reported that more than 100 phytocannabinoids are isolated and characterized to have active interaction with the endocannabinoid system ^8^. Amongst these phytocannabinoids, Δ^9^-tetrahydrocannabinol (THC) and cannabidiol (CBD) are the most well-studied. Being a psychoactive constituent of *Cannabis sativa*, the use of THC as a pharmacological therapy is therefore limited, which draws further attention towards non-psychoactive phytocannabinoids. The non-psychoactive constituent CBD has shown to possess protective properties in various diseases including seizure, cancer and food disorders ^9-11^. As a neuroprotective agent, CBD has been successfully approved for treatment of epilepsy by the US Food and Drug Administration (FDA) in 2018. This has opened up avenues to further study the therapeutic values of other non-psychoactive phytocannabinoids including cannabidivarin (CBDV), cannabichromene (CBC) and cannabigerol (CBG) ^12^. N-arachidonoylethanolamine (AEA) and 2-arachidonoylglycerol (2-AG) are the most important endocannabinoids in the canonical endocannabinoid system, and many studies have documented their roles in mediating physiological processes, such as neuroinflammation, pain perception and metabolism, through binding to CB1/2 receptors in mammals ^6,11^. In addition, numerous synthetic cannabinoids are constructed as agonists/antagonists of CB1/2 receptors, and further utilized to probe into the physiological functions of the endocannabinoid system ^13^. Although CB1/2 receptors are known to modulate the effects of these cannabinoids, non-canonical G protein-coupled receptors (GPCRs), such as transient receptor potential vanilloid (TRPV) channels, peroxisome proliferator-activated receptor (PPAR), orphan GPCR GPR55 and 5-HT1A, have also been reported to be pharmacological targets of these cannabinoids in mediating various physiological functions ^11^.

The fruit fly, *Drosophila melanogaster*, is an excellent model as a platform for screening new drugs and unraveling molecular mechanisms ^14^. Due to its conserved genome and sophisticated genetic tools, it has been widely used to study biological processes and diseases, such as obesity, neurological disorders and longevity. Despite the lack of canonical cannabinoid CB1/2 receptors, various orthologs of endocannabinoid-metabolized enzymes and non-canonical receptors are well conserved in *Drosophila* ^15,16^. The major endocannabinoids, namely AEA and 2-AG, are expressed at low levels in *Drosophila* larvae as reported by some labs ^17,18^, although undetectable in other studies ^19,20^. Nevertheless, the endocannabinoid-like signal lipid 2-linoleoyl glycerol (2-LG) is abundant in flies ^16,20^. These raise the possibility that cannabinoids may impact on the physiology and behaviors of *Drosophila*. Interestingly, a few recent studies have highlighted potential therapeutic effects of cannabinoids in *Drosophila* for treating various human diseases including cardiovascular diseases, seizures and Parkinson’s disease despite the absence of a canonical cannabinoid-signaling pathway ^21-23^. Inhalation of vaporized marijuana leads to an increase in cardiac contractility in adult flies ^21^. Importantly, the neuroprotective effects of cannabinoids are evident with exogenous application of AEA and 2-AG conferring neuroprotection against seizures in multiple fly models ^22^. Pharmacological treatment with the synthetic cannabinoid CP55940, a non-selective CB1/2 receptor agonist, prolongs survival and restores locomotor activity against paraquat toxicity in flies ^23^. Thus, *Drosophila* is emerging as a valuable model to elucidate the functions and pathways mediated by cannabinoids in a CB1/2 receptors-independent manner. However, little is known on the role of cannabinoids in food intake and metabolism in *Drosophila*. Studies using rodent models of feeding disorders have demonstrated that treatment with endocannabinoids positively regulates food consumption ^24-26^. In contrast, exogenous AEA and 2-AG significantly inhibit food intake in *C. elegans* ^27^. With the discrepancy in findings using different models, more research is required to determine the role of cannabinoids in food intake.

Here, we report cannabinoid preference and its inhibitory impact on food intake in *Drosophila*. We show that adult flies have an innate ability to detect and develop a preference for food containing cannabinoids, independent of their sensory inputs. Surprisingly, treatment with phyto- and endo-cannabinoids leads to a decrease in food intake. Interestingly, endocannabinoid-induced suppression in food intake is attenuated by AM251, a non-selective CB1 antagonist. Furthermore, we show that the endocannabinoid AEA prolongs starvation resistance and reduces lipid metabolism. Overall, this is the first systematic study demonstrating cannabinoid preference and its impact on feeding behavior in *Drosophila*.

## Results

### Flies prefer to consume food containing phytocannabinoids

The non-psychoactive phytocannabinoids are naturally occurring compounds derived from *Cannabis sativa*, and exert their biological effects via interaction with cannabinoid receptors or other G-protein coupled receptors ^28^. To investigate any potential role of cannabinoids in food preference in adult flies, we took advantage of the two-choice CAFE assay ^29^, in which flies are presented with cannabinoid food and proper control food in four capillaries (Figure 1A). We first assessed whether flies display a preference for four phytocannabinoids, including CBD, CBDV, CBC and CBG. Flies were presented with various concentrations of CBD (0.01 mg/ml, 0.1 mg/ml or 1 mg/ml), CBDV, CBC or CBG (0.01 mg/ml or 0.1 mg/ml) in each CAFE assay. Statistical analysis shows that the flies did not exhibit obvious preference for food with the individual phytocannabinoids in the first 2 days (Figures 1B-E). Interestingly, a significant preference for these phytocannabinoids was observed in the following two days. Flies showed the preference for consuming food with 0.1 mg/ml or 1 mg/ml CBD on day 3 and day 4, however, appeared to prefer 0.01 mg/ml CBD only on day 4 (Figure 1B). These data indicate that flies possess an ability to detect CBD and develop a dose-dependent CBD preference over time. Likewise, food containing high concentration of CBDV (Figure 1C), CBC (Figure 1D) or CBG (Figure 1E), was preferentially consumed by the flies on day 3 and day 4. Flies also displayed food preference to CBG at the lower concentration on day 4 (Figure 1E), similar to CBD (Figure 1B). Thus, these data indicate that flies can develop a delayed preference for phytocannabinoids in a dose-dependent manner.

**Figure 1.**
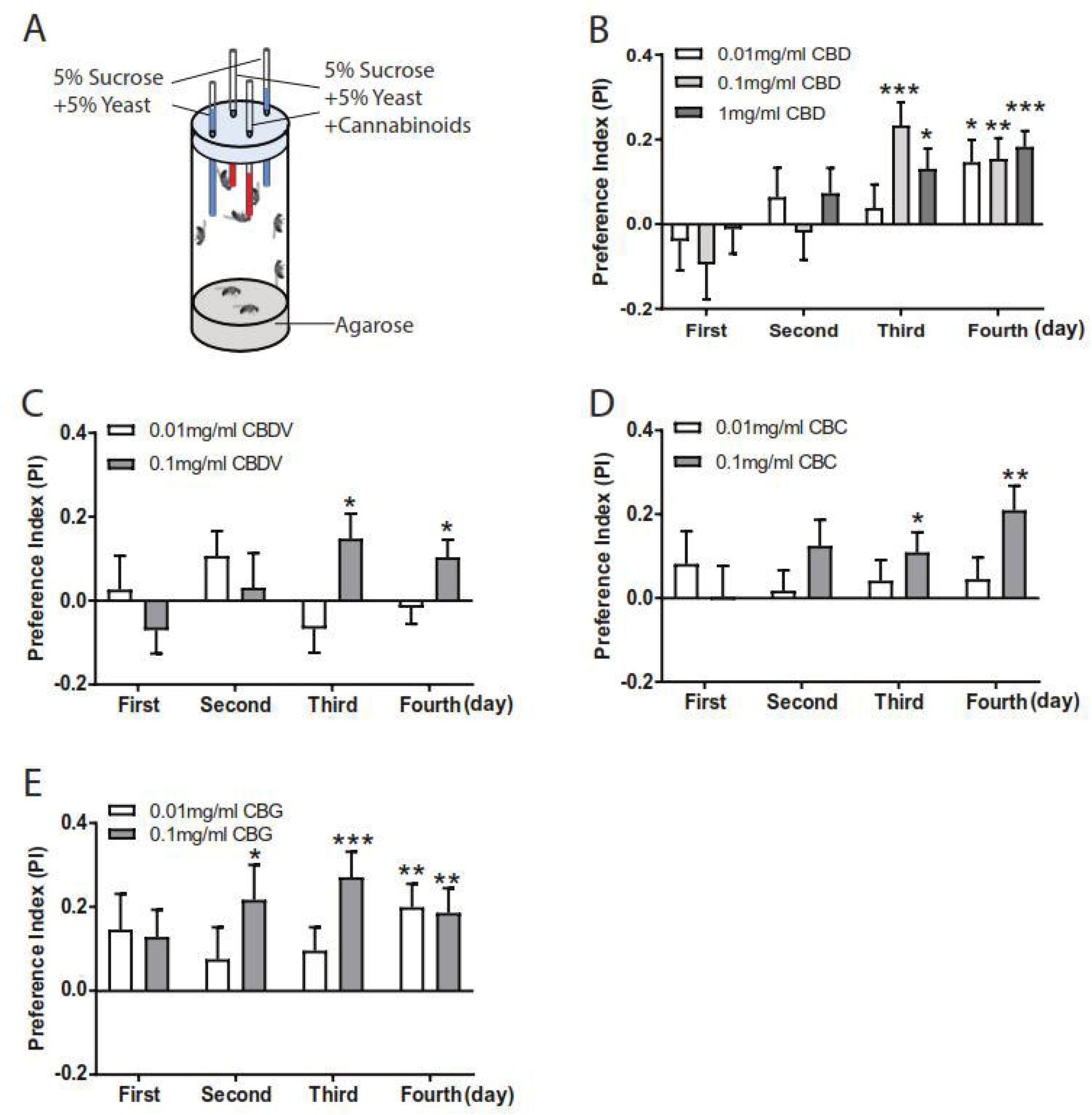
Flies developed a preference for phytocannabinoids. (**A**) Schematic diagram of the CAFE assay for examining cannabinoid preference. Eight flies in a single vial were presented with four capillaries in which two contained cannabinoids and the other two with normal liquid food for four consecutive days. (**B-E**) Flies exhibited and developed preference for phytocannabinoids in a time-dependent manner. (**B**) Delayed preference for 0.1 mg/ml and 1 mg/ml CBD was detected on day 3 and day 4. A significant increase in preference for food containing 0.01 mg/ml CBD was observed on day 4. Flies displayed preference for food containing 0.1 mg/ml CBDV (**C**) and CBC (**D**) on day 3 and day 4. Similar to CBD, the preference for CBG-containing liquid food was significantly increased from day 2 to day 4 (n=17-28 vials/group). Data are represented as mean ± S.E.M. One-sample t-test was applied to determine statistical significance. \**p*<0.05, \*\**p*<0.01, and \*\*\**p*<0.001.

### Flies prefer to consume food with endocannabinoids and synthetic cannabinoids

We next extended the characterization of food preference to endocannabinoids and synthetic cannabinoids. Flies were presented with either of endocannabinoids, namely anandamide (AEA) and 2-arachidonoylglycerol (2-AG), at 0.01 mg/ml and 0.1 mg/ml (Figure 2A and 2B). The preference indexes of food intake indicate a robust preference for food with a high concentration of AEA (Figure 2A) and 2-AG (Figure 2B), but not with the lower concentration. The strong preference for 2-AG was sustained throughout 4 days of feeding (Figure 2B), whereas the preference for AEA remained consistent, albeit relatively lower, in the first 3 days (Figure 2A). Previous studies have reported the identification of the endocannabinoid-like signal lipid 2-LG, an evolutionary alternative to 2-AG in *Drosophila* ^16,20^. Arachidonic acid (AA) supplement can promote the synthesis of 2-AG in *Drosophila* ^16^. We also observed a weak but significant preference for food containing 2-LG only on day 3, in contrast to more robust preference for food with AA throughout the entire course of feeding (Figure 2C).

**Figure 2.**
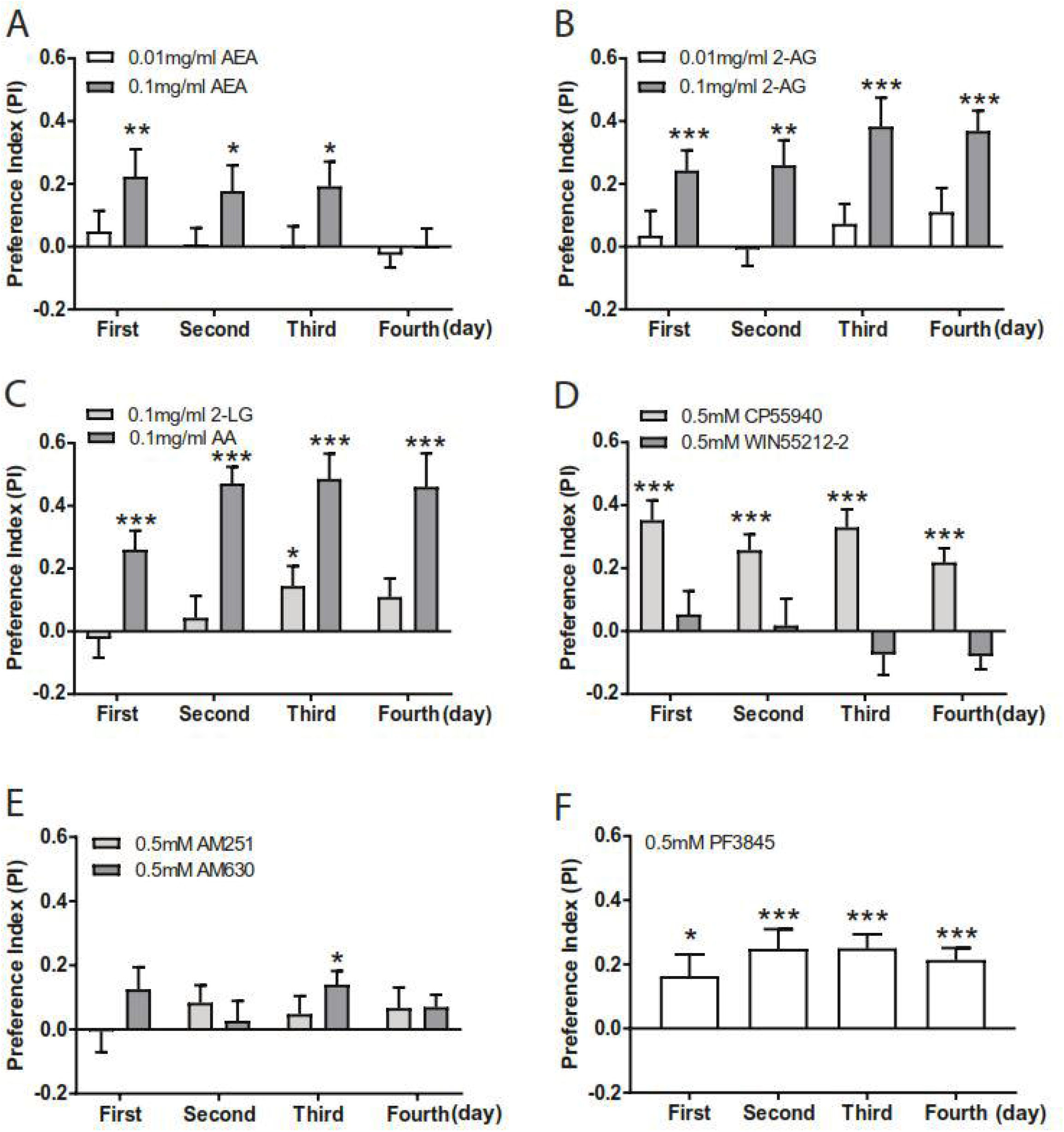
Flies preferentially consumed endocannabinoids and synthetic cannabinoids. Flies were presented with food containing endocannabinoids and synthetic cannabinoids for four consecutive days. Flies preferentially consumed food with endocannabinoids, including 0.1 mg/ml AEA (**A**), 0.1 mg/ml 2-AG (**B**) and 0.1 mg/ml AA (n=18-21). However, a significant preference for 0.1 mg/ml 2-LG was detected only on day 3 (**C**). (**D**) A selective preference for 0.5 mM CB1/2 receptors agonist CP55940, but not WIN55212-2, was detected for four days (n=18-20). (**E**) Flies did not consistently display significant preference for 0.5 mM CB1/2 receptors antagonists AM251 or AM630 (n=18-24). (**F**) Flies displayed a strong preference for 0.5 mM PF3845 over the entire feeding course (n=21). Data are mean ± S.E.M. One-sample t-test was applied to determine statistical significance. \**p*<0.05, \*\**p*<0.01, and \*\*\**p*<0.001.

With the detection of food preference for endocannabinoids in flies, we further assessed potential preference for synthetic cannabinoids. Flies exhibited a significant preference for food containing 0.5 mM CP55940 over 4 days (Figure 2D), but not for another CB1/CB2 receptor agonist WIN55212-2 at the same concentration (Figure 2D). Given the absence of CB1/2 receptors in the fly genome, the results suggest that these two synthetic cannabinoids might target distinct receptors in Drosophila. Furthermore, we also investigated potential effect on food preference for AM251 (0.5 mM) or AM630 (0.5 mM), two antagonists of CB1 and CB2 receptors, respectively. The consumption for AM251 or AM630-containing food remained similar to that for the control food throughout 4 days of feeding, despite a weak preference for AM630 on day 3 (Figure 2E). Fatty acid amide hydrolase (FAAH), which is well conserved between mammals and *Drosophila* ^30^, catabolizes AEA into AA and ethanolamine ^31^. Treatment with PF3845, an inhibitor of mammalian FAAH, has also been shown to increase the endocannabinoids in fly hemolymph ^30^. Flies displayed a significant preference for food with 0.5 mM PF3845, and the preference index remained relatively high throughout 4 days of feeding (Figure 2F). Taken together, these findings indicate a general preference for both endocannabinoids (AEA and 2-AG) and synthetic cannabinoid CP55940 in flies.

### Cannabinoid preference is independent of sensory inputs in flies

To examine whether the preferential response to cannabinoids is due to sensory stimuli in flies, we examined possible involvement of gustatory and olfactory functions in food preference using taste-deficiency mutants (*poxn*^70-23^ and *poxn* ^Δ^^M22-B5^) (Figure 3A and 3B), and olfaction-deficiency mutants (*orco*^1^ and *orco*^2^) (Figure 3C and 3D), respectively. In comparison to the control *w*^1118^ flies, both *poxn* and *orco* homozygotes displayed similar preference for 1 mg/ml CBD and 0.5 mM CP55940 over 4 consecutive days of feeding, suggesting that the preference for cannabinoids in flies is modulated through a mechanism independent of gustatory and olfactory sensory functions.

**Figure 3.**
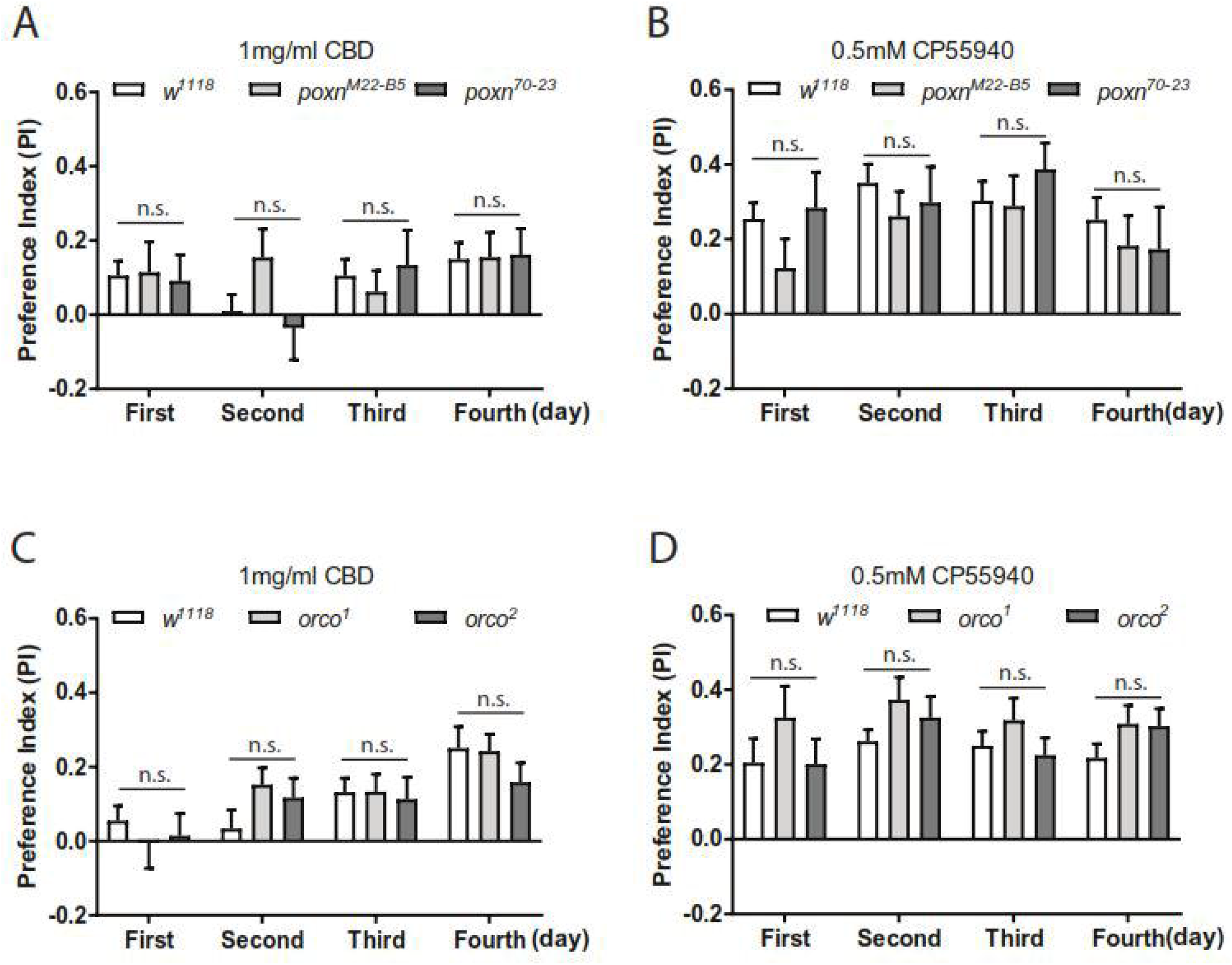
The preference for cannabinoids is independent of gustatory and olfactory inputs. The preference for 1 mg/ml CBD and 0.5 mM CP55940 was measured in two *poxn* mutants (**A, B**) and *orco* mutants (**C, D**) over four consecutive days. Homozygous *poxn*^ΔM22-B5^ and *poxn*^70-23^ mutants, and *orco*^1^ and *orco*^2^ mutants exhibited similar preference for CBD (**A, C**) and CP55940 (**B, D**) as compared to the control *w*^1118^ flies (n=11-24 vials/ group). Data are represented as mean ± S.E.M. One-way ANOVA followed by Dunnett’s post-hoc test. n.s., not significant.

### An inhibitory effect of phytocannabinoids on food intake

The lack of gustatory and olfactory sensory inputs in influencing cannabinoid preference suggests a potential pharmacological effect of these compounds on metabolism in *Drosophila*. Phytocannabinoids have been shown to affect food intake, and can be developed as potential therapeutic agents for obesity treatment ^12^. We next investigated if phytocannabinoids regulate food intake in flies. The consumption of normal food by flies was quantified for two consecutive days following two days of cannabinoid training (Figure 4A). Pre-treatment of the flies with higher concentrations of CBD at 1 mg/ml and 2 mg/ml, but not with the low concentration at 0.1 mg/ml, significantly decreased the total food intake on both days (Figure 4B). The movement ability of flies was assessed to examine whether the effect observed in food intake was due to a decline in locomotion. In the negative geotaxis assay, the treatment with high concentration of CBD did not impair the climbing ability of adult flies (Figure S1A), suggesting that decreased food intake by CBD treatment is not attributed to altered locomotion. However, food intake was significantly decreased only on day 2 following the treatment with 0.1 mg/ml CBC and CBDV (Figure 4C). No or negligible effect was detectable upon CBG treatment in flies (Figure 4C). Subtle effects of CBC, CBDV and CBG could be due to their lower feeding concentration. We were unable to further increase the feeding concentration, given the low concentration of commercially available stock solutions. During the first two days of training, the amounts of food containing phytocannabinoids or the respective control solutions consumed by the flies remained similar (Figure S2). These control experiments suggest that the decrease in food intake is not caused by altered food consumption during the initial cannabinoid training. Thus, our results indicate that phytocannabinoids elicit a functional effect on the regulation of food intake in flies.

**Figure 4.**
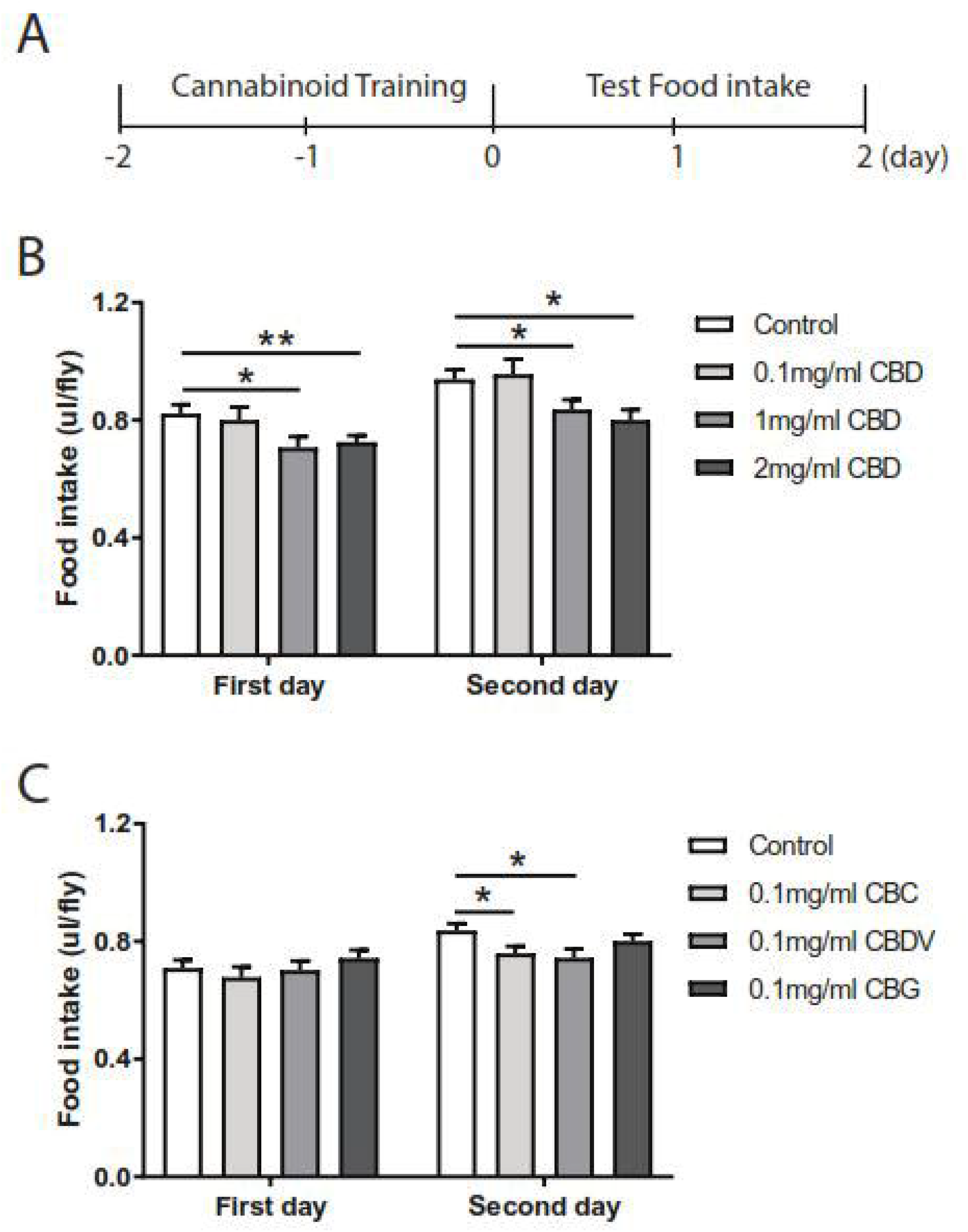
Phytocannabinoids reduce food intake in flies. (**A**) A diagram of the experimental scheme for the food intake assays. Following two days of cannabinoid treatment (day -2 and day -1), the normal food intake by the flies was measured for two consecutive days (day 1 and day 2) as depicted in the histograms (**B, C**). (**B**) CBD at 1mg/ml and 2 mg/ml, but not at 0.1 mg/ml, significantly decreased normal food intake on both day 1 and day 2 (n=13-19 vials/group). (**C**) Pre-treatment with 0.1 mg/ml CBC and CBDV, but not with CBG, led to significant reductions in food intake on day 2 (n=32-33 vials/group). All data are represented as mean ± S.E.M. One-way ANOVA followed by Dunnett’s post-hoc test. \**p*<0.05 and \*\**p*<0.01.

### Endocannabinoids inhibit food intake

Growing studies have reported that endocannabinoid signaling functions to enhance food intake through the activation of CB1 receptor in mammals ^25^. However, the sequence homology analysis indicated that canonical CB1 receptor appears to be absent in *Drosophila* ^15,32^. Cannabinoids might function in an alternative way in flies, for example, via non-canonical cannabinoid receptors. To explore whether the endocannabinoids regulate food consumption, flies were pre-treated with AEA and 2-AG at the concentration of 0.01, 0.1 and 0.5 mg/ml in the first two days prior to the measurement of food intake in the next two days (Figure 4A). The pre-treatment with higher concentrations of AEA exhibited profound effects on food intake whereby consumption of normal food was significantly attenuated on both days (Figure 5A). Consistently, a significant reduction in food intake was observed with 2-AG treatment at the higher concentrations (Figure 5B). However, this effect was only observed on the first day following the pre-treatment. Strikingly, the amounts of food containing the higher concentrations of AEA or 2-AG consumed by flies were significantly lower as compared to those of the normal food (Figure S3A and S3B). As a control, the climbing ability was not affected by either AEA or 2-AG treatment (Figure S1B and S1C). Next, we attempted to determine the possible effect of 2-LG (0.01 and 0.1 mg/ml) on food intake. Similar to 2-AG treatment, flies consumed significantly less food with 2-LG at the higher concentration of 0.1 mg/ml as compared to the normal food (Figure S3C) during the initial training. The pre-treatment with higher concentration of 2-LG also significantly decreased food consumption in the next two days (Figure 5C).

**Figure 5.**
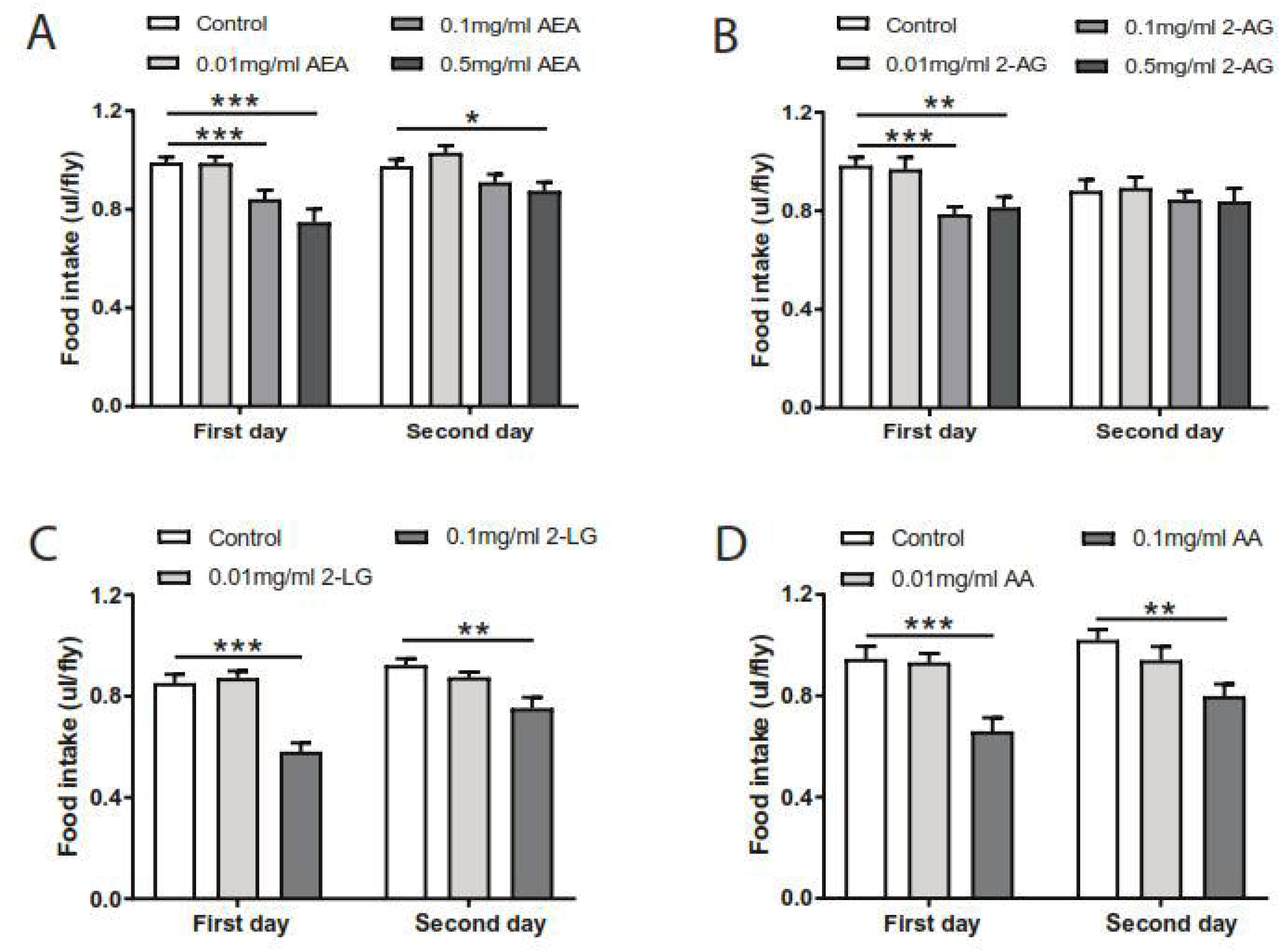
Endocannabinoids decrease food intake in flies. Pre-treatment with AEA (**A**) and 2-AG (**B**) led to significant reductions in food consumption on day 1, with a weak effect of AEA on day 2 (n=12-27). More profound effects of 2-LG (**C**) and AA (**D**) were detected. The food intake by the flies was significantly decreased with 0.1 mg/ml 2-LG (**C**) and AA (**D**) on both day 1 and day 2 (n=16-18). Data are represented as mean ± S.E.M. One-way ANOVA followed by Dunnett’s post-hoc test. \**p*<0.05, \*\**p*<0.01, and \*\*\**p*<0.001.

A recent study has reported that AA facilitates 2-AG synthesis by the fly ortholog of diacylglycerol lipase dDAGL ^16^. Since both 2-AG and 2-LG account for the reduction of food intake, we postulated that treatment with AA in flies may also have an inhibitory role in food intake. In agreement with our hypothesis, a similar inhibitory effect on food consumption was observed upon AA treatment. Normal food consumed by flies treated with higher concentration of AA at 0.1 mg/ml was significantly reduced as compared to the control group (Figure 5D). Similarly, food consumption was remarkably lower during the first 2 days of 0.1 mg/ml AA pre-treatment (Figure S3D), and the climbing ability was not affected by AA treatment (Figure S1D). Together, these results suggest that endocannabinoids, like phytocannabinoids, elicit an inhibitory effect on the regulation of food consumption.

### AM251 attenuated the inhibitory effect of AEA on food intake

It has been documented that the activity and levels of AEA are negatively regulated via at least two models. First, AEA is degraded by FAAH to AA and ethanolamine ^31^. Second, fatty acid-binding protein (FABP) facilitates the transportation of AEA to FAAH, which leads to subsequent degradation and inactivation of AEA ^33^. Overexpression of FABP significantly promotes AEA uptake and hydrolysis in neuroblastoma cells ^34^. Given that AEA treatment led to a reduction in food intake, we next examined whether changes in endogenous AEA levels through FAAH inhibition or FABP overexpression can affect food intake. Importantly, treatment with the FAAH inhibitor PF3845, which inhibit endocannabinoid degradation in flies ^30^, led to a significant reduction in food intake on day 1 (Figure 6A). Likewise, food consumption was significantly reduced during PF3845 pre-treatment (Figure S3E). Conversely, overexpression of fly dFABP significantly enhanced the food intake on two consecutive days (Figure 6B). These results are in line with an inhibitory effect of AEA on food intake.

**Figure 6.**
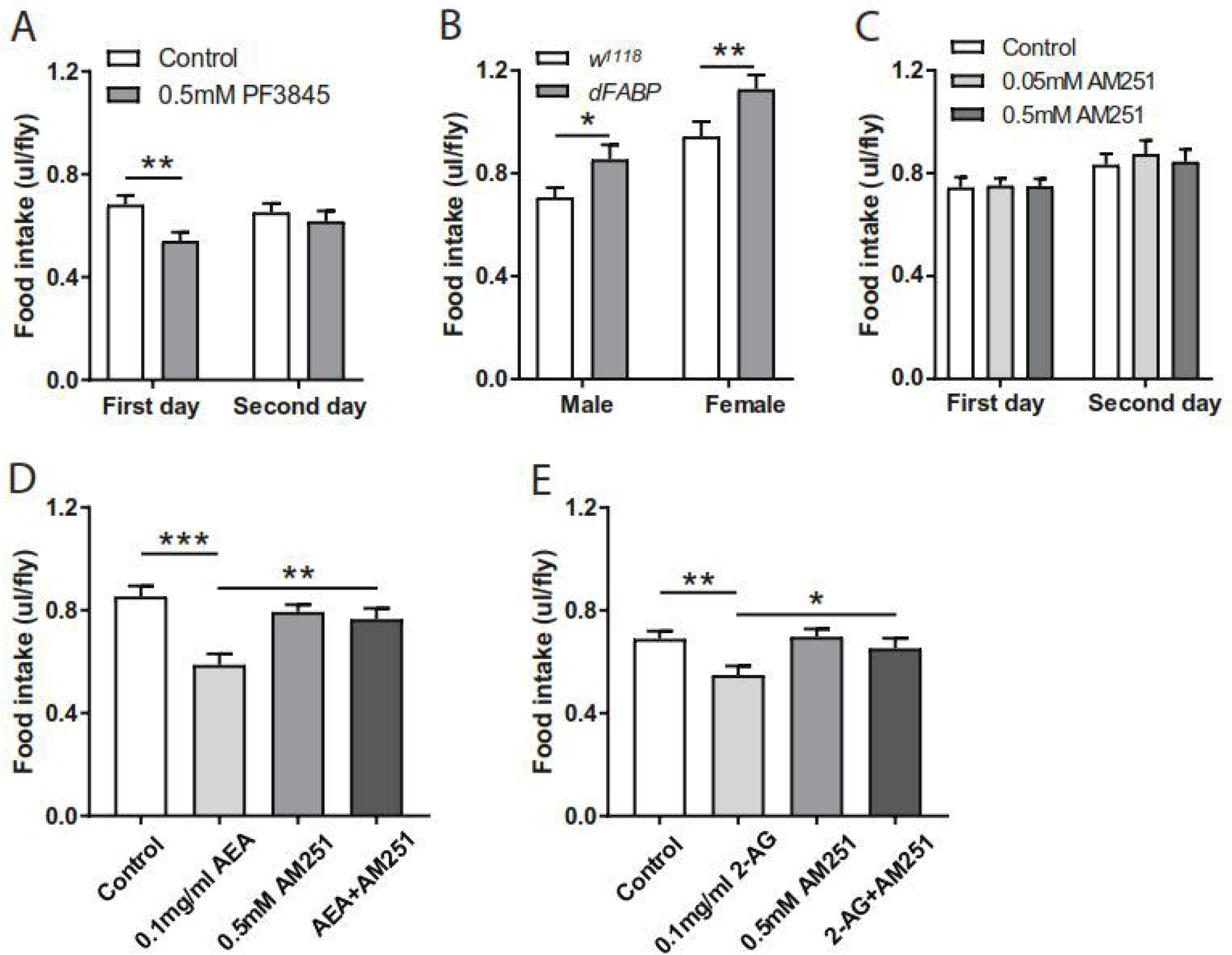
AM251 attenuated the inhibitory effects of AEA on food intake. (**A**) While inhibition of FAAH with 0.5 mM PF3845 led to a significant decrease in the consumption of normal food on day 1 (n=18). (**B**) Overexpression of dFABP in flies significantly increased food intake (n=16-19). (**C**) Food intake was not altered following AM251 pre-treatment (n=18). Administration of 0.5 mM AM251 significantly mitigated the decreases in food intake induced by 0.1 mg/ml AEA (**D**) and 0.1 mg/ml 2-AG (**F**) on day 1 (n=16-24 vials/group). All data are represented as mean ± S.E.M. One-sample t-test and one-way ANOVA followed by Dunnett’s post-hoc test were used to analyze the difference. \**p*<0.05, \*\**p*<0.01, and ***p<0.001.

Next, we sought to determine whether the CB1 receptor antagonist AM251 (0.5 mM) can counteract AEA (0.1 mg/ml) or 2-AG (0.1 mg/ml) effect in terms of food intake. Flies pre-fed with AM251 did not alter the levels of food intake in the following two days (Figure 6C). Flies also consumed similar amount of food containing AM251 during the pre-treatment (Figure S3F). These results indicate that the treatment with this CB1 receptor antagonist did not affect food consumption in flies. Treatment with 0.1 mg/ml AEA or 2-AG led to significant reductions in normal food consumption (Figures 6D and 6E). Interestingly, co-treatment of AM251 with AEA (Figure 6D) or 2-AG (Figure 6E) significantly attenuated their inhibitory effects on food intake. Due to the lack of canonical CB1/2 receptors in *Drosophila* genome, AEA and 2-AG regulate food intake, likely via another unknown receptor that can be blocked by AM251.

### AEA treatment promotes starvation resistance and inhibits lipid metabolism

Given that phyto- and endo-cannabinoids can modulate food intake in flies, we next tested whether they also affect starvation resistance and lipid metabolism. Starvation resistance was quantified by measuring the survival percentage of the flies following cannabinoid treatment. After treatment with 0.1 mg/ml AEA, 0.1 mg/ml AA or 0.5 mM CP55940 for 2 days, flies exhibited significant increases in the survival rate (Figures 7A, 7C and 7E), suggesting an enhanced resistance to starvation. Unexpectedly, neither 2-AG nor CBD pre-treatment showed any effect on starvation resistance (Figures 7B and 7D). The observation of enhanced starvation resistance further prompted us to investigate whether body fat deposition was affected by AEA. Following 18 h starvation, the levels of triglyceride (TAG) were measured in endocannabinoid-treated flies. TAG levels in AEA- or AA-treated flies were significantly elevated when compared to the control flies (Figure 7F). In line with no effect on starvation resistance, we did not observe a significant alteration in TAG levels upon 2-AG treatment (Figure 7F). Thus, AEA and AA can enhance the resistance to starvation likely through the inhibition of lipid metabolism.

**Figure 7.**
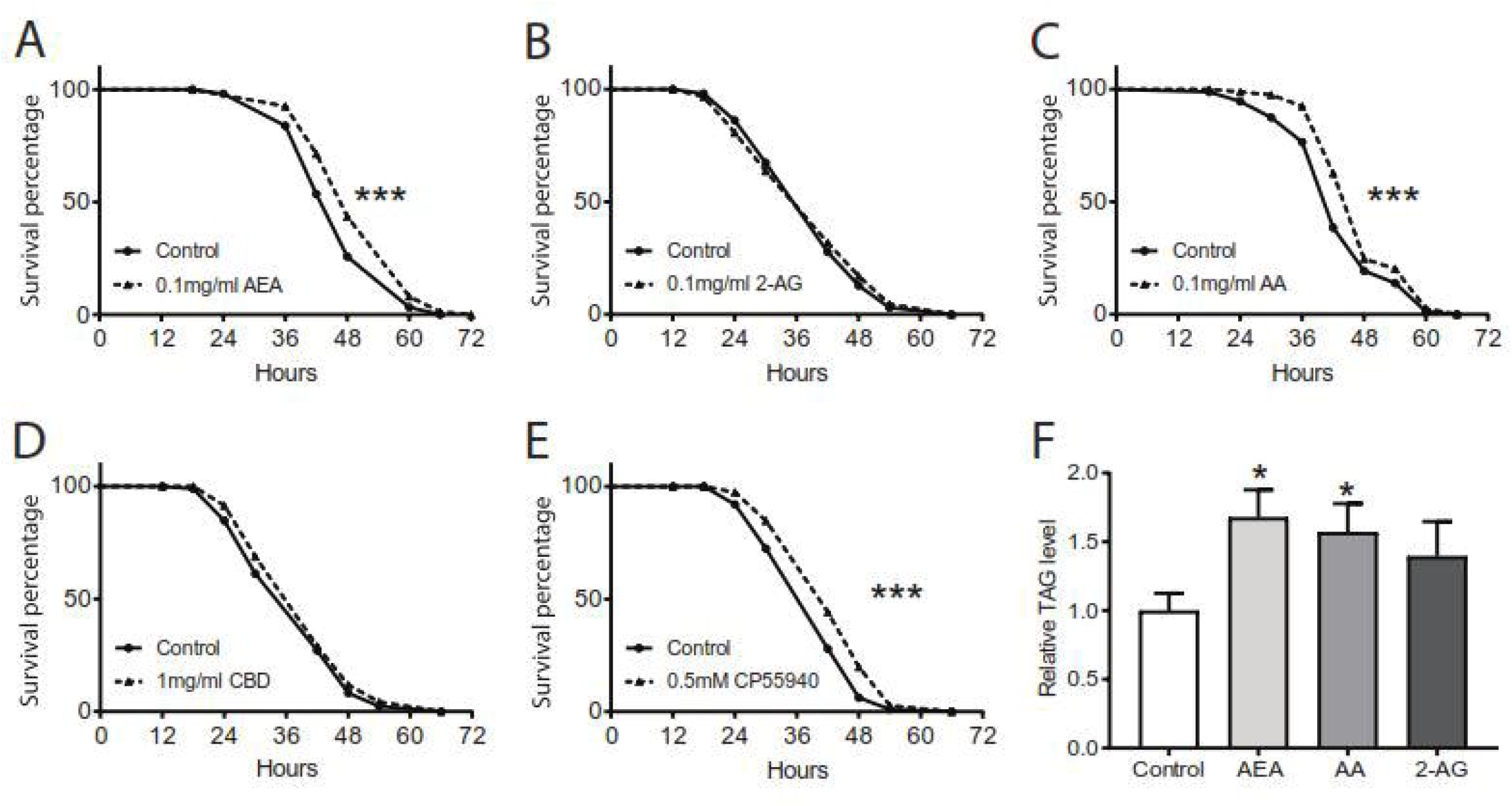
AEA and AA protects flies from starvation stress and inhibits lipid metabolism. (**A-E**) Following two days of cannabinoid treatment, the percentage of the surviving flies was quantified to assess the resistance to starvation. Pre-treatment with 0.1 mg/ml AEA (**A**), 0.1 mg/ml AA (**C**), and 0.5 mM CP55940 (**E**) significantly enhanced the starvation resistance, while no effect was detected with 0.1 mg/ml 2-AG (**B**) and 1 mg/ml CBD (**D**) (n=9-14 vials/group). (**F**) Quantification of the triacylglycerol (TAG) levels in the flies revealed a significant increase with 0.1 mg/ml AEA or 0.1 mg/ml AA pre-treatment following 18 h starvation as compared to the control group (n=7-10 vials/group). Both one-sample t-test and Log-rank (Mantel-Cox) test were used to determine the significant differences. \**p*<0.05 and \*\*\**p*<0.001.

## Discussion

*Drosophila* has been developed as an important invertebrate model for drug screening due to its conserved genome and biological machineries. In this study, we utilized this model to investigate the effects of a spectrum of cannabinoids including phytocannabinoids, endocannabinoids and synthetic cannabinoids on food preference and consumption. Our findings reveal cannabinoid preference as well as an inhibitory function of cannabinoids in food intake in flies. As canonical CB1/2 receptors are absent in flies, this study provides insight into the pharmacological role of cannabinoids in the modulation of food consumption via a non-canonical cannabinoid signaling pathway.

*Cannabis* extracts are known to possess pesticidal properties, which can effectively repel insects and inhibit the growth of microbial pathogens ^35^. While the naturally occurring phytocannabinoids display potential pharmacological roles, it is still poorly understood whether *Drosophila* is a useful model for understanding the functions of phytocannabinoids. Here, we characterize the preference for consuming cannabinoids in flies by using the two-choice feeding assay. The results indicate that flies display a significant preference for phytocannabinoids over time. A recent study has showed that Tobacco Hornworm *Manduca sexta* larvae prefer the food with low concentration of CBD over higher dose of CBD, and high CBD concentration is detrimental for larvae development ^36^. However, no aversive response to phytocannabinoids was detected in this study. One possibility is that relatively low concentrations of phytocannabinoids were administrated in our CAFE assays. While we also observed a delayed development of fly larvae upon CBD treatment (data not shown), our current findings indicate these phytocannabinoids are not toxic to adult flies, at least at the relatively low concentrations used in this study.

In contrast to phytocannabinoids, flies displayed quicker and stronger preference for endocannabinoids and psychoactive cannabinoid CP55940. Although endocannabinoids are present in various vertebrates, very low amount of AEA and 2-AG is produced in flies ^17,18^. AA, a critical substrate for eicosanoid biosynthesis, is also almost undetectable in flies ^37^. A previous behavioral study in *Drosophila* has revealed a stage-specific discrimination of fatty acids. Larvae prefer various unsaturated fatty acids, but adult flies develop a preference for the diet containing high saturated fatty acids ^38^. In this present study, we found that no apparent toxicity and aversive responses to these cannabinoids were detectable in flies, consistently, sensory inputs from gustatory and olfactory systems did not influence cannabinoid preference. These findings suggest that the cannabinoids may have some pharmacological roles in *Drosophila*. Several previous studies have highlighted the protective effects of the endocannabinoids and the synthetic cannabinoid CP55940 through non-canonical signaling pathways in *Drosophila*. Endocannabinoids and their metabolite AA were shown to possess anti-convulsant properties in protecting against seizures through the TRP channel Water witch in fly seizure models ^22^; CP55940 alleviates paraquat-induced toxicity with the observations of increased survival and locomotion via the JNK signaling pathway ^23^. Thus, our studies and others suggest that *Drosophila* may be an alternative model to dissect CB1/2 receptors-independent roles of cannabinoids as well as their pharmacological functions in diseases.

To understand whether cannabinoids affect food intake and metabolism, we systematically characterized the effects of various available cannabinoids in flies. The food intake was found to significantly decrease following pre-treatment with CBD, which is consistent with previous mammalian studies illustrating the effectiveness of CBD in controlling food consumption and preventing obesity in rodents ^11,39,40^. Although these results in both mammals and flies are seemingly similar, the underlying mechanisms apparently differ. CBD inhibits food intake by blocking CB1 receptor and activating CB2 receptor in rodents ^11^. By contrast, in flies lacking CB1/2 receptors, CBD functions through a CB1/2 receptor-independent mechanism. Our study is the first to illustrate the pharmacological function of CBDV and CBC whereby they can reduce food consumption. However, CBG does not seem to affect the food intake in *Drosophila*, whereas in rats CBG promotes food consumption at a high concentration ^41^. The lack of CBG effect on food intake in flies could be due to the administration of CBG at a lower concentration in this study. Overall, these phytocannabinoids did not display a strong phenotype, thus, their potential as a treatment for eating disorders requires further interrogations. Although phytocannabinoids can bind to CB1/2 receptors, they display higher affinity for other GPCRs, such as GPR55, TPRV channels and PPARγ ^42,43^. Since these non-canonical cannabinoid receptors are well conserved, it is conceivable that phytocannabinoids may exert their protective roles through modulation of these receptors in flies. Thus, further studies are required to investigate whether and how phytocannabinoids regulate food intake via one of these GPCRs.

Similar to the actions of phytocannabinoids, the present study puts forward the idea that the endocannabinoid-mediated pathway may modulate food intake. Pre-treatment with AEA and 2-AG significantly inhibited the food intake in adult flies, which is consistent with the finding in *C. elegans* ^27^. However, administration of AEA and 2-AG promotes food intake and increase odor detection via binding to CB1 receptor in rodents ^26,44^. As there is a lack of cannabinoid receptors to transduce AEA and 2-AG signals in *Drosophila* ^15,32^, they may function through pathways central to other lipids of N-acyl amides and 2-acyl glycerols. Although endocannabinoid-mediated downstream pathway in food intake remains to be determined, our findings reveal that 2-LG, a closely-related 2-acyl glycerol ligand, also shows inhibition of food intake, suggesting a possible existence of a signaling pathway for the regulation of food intake in *Drosophila*. Furthermore, previous studies have also reported that OEA reduced food intake in rodents and goldfish ^45,46^. Both OEA and AEA belong to a family of fatty acid ethanolamides (FAEs). It is conceivable that endocannabioids in *Drosophila* regulates food intake via a mechanism, likely similar to other lipids of N-acyl amides and 2-acyl glycerols. Moreover, we also found that flies pre-treated with FAAH inhibitor PF3845 consumed less food, whereas overexpressing dFABP in flies increased the food intake. These data suggest that FAAH-induced AEA metabolism and FABP-mediated AEA inactivation are important for this feeding behavior, thus reinforcing the crucial role of AEA in regulating food intake. Interestingly, the CB1 receptor antagonist, AM251, counteracts the inhibitory effects of AEA and 2-AG on food intake. This could possibly be due to the interaction of endocannabinoids with known non-CB receptors or other unidentified CB1-like receptors which can be targeted by AM251. In addition to its primary role as a CB1/2 receptor antagonist, AM251 is found to facilitate the activation of GPR55 that is a known target of AEA and AA ^47,48^. From our findings, we speculate that AEA and 2-AG might regulate food intake via a GPR55 ortholog in *Drosophila*. To date, no fly GPR55 ortholog has been characterized. Further work is required to identify a possible GPR55 ortholog and its implications in food intake in flies.

The inhibitory role of cannabinoids in food preference and intake is particularly interesting as there is an absence of CB1/2 receptors in *Drosophila*. First, phytocannabinoids have been shown to exhibit high affinity for various non-CB1/2 receptors, such as GPR55, TPR channels and PPARγ ^42,43^, whereas the binding affinity to non-CB1/2 receptors is comparatively lower than CB1/2 receptors for endocannabinoids ^11^. The extent of cannabinoid preference and cannabinoid-induced inhibition of food intake could be due to differential binding affinities to the receptors in flies. It is known that the structure of these cannabinoids determines its functional activity through the binding affinity to the CB1/2 receptors ^49^. However, it is unknown how the structural element can affect the binding state of these cannabinoids to the non-canonical receptors. Second, the inhibitory effect of AEA and its metabolite AA could possibly be due to the decrease in lipid metabolism. It is very likely that the increase in TAG levels could underlie the enhancement of starvation resistance. Although 2-AG exhibited a similar outcome in food intake to AEA, it is surprising that 2-AG did not affect lipid metabolism. Possible mechanism of AEA action in lipid metabolism awaits future investigations.

In conclusion, our study demonstrates that adult flies can detect and develop the preference for phyto-, endo- and synthetic cannabinoids. Importantly, these cannabinoids can inhibit food intake. We show that the endocannabinoids (AEA and 2-AG) inhibit the food intake via an AM251-targeted pathway. Moreover, the endocannabinoid AEA modulates food intake and starvation resistance probably through low lipid metabolism.

## Methods

### Fly stocks and maintenance

The following fly stocks were obtained from Bloomington *Drosophila* stock center (BDSC; Indiana, USA): Canton S (#64349), *w*^1118^ (#5905), *orco*^1^ (#23129), *orco*^2^ (#23130), and *poxn*^70-23^ (#60688). The following fly stocks were kindly provided: *poxn* ^Δ^^M22-B5^ (M. Noll from University of Zurich, Switzerland) ^50^, *w*^1118^ (isoCJ1) and *dFABP* (J.C.P. Yin from University of Wisconscin-Madison, USA) ^51^. Flies were reared on standard medium at 25°C and approximately 70% humidity under a 12 h light/12 h dark cycle. 3-5-day-old adult male flies were collected using light CO_2_ anesthesia and allowed to recover for 2 days before further experimentation.

### Drug selection and delivery

Various cannabinoids used in this study were categorized into 1) phytocannabinoids, 2) endocannabinoids, and 3) synthetic cannabinoids (Table 1). These drugs were dissolved in their respective solvents and further diluted to the desired concentrations in liquid food (5% sucrose and 5% yeast extract).

**Table 1.** Phytocannabinoids, endocannabinoids and synthetic cannabinoids used in this study.

| Cannabinoids | Name | Stock | Solvent | Source |
| --- | --- | --- | --- | --- |
| Phytocannabinoids | CBD | 25 mg/ml | Ethanol | Cayman Chemicals |
|  | CBDV | 1 mg/ml | Methanol | Cayman Chemicals |
|  | CBC | 1 mg/ml | Methanol | Cayman Chemicals |
|  | CBG | 1 mg/ml | Methanol | Cayman Chemicals |
| Endocannabinoids | AEA | 5 mg/ml | Ethanol | Tocris Bioscience |
|  | 2-AG | 5 mg/ml | Ethanol | Tocris Bioscience |
|  | 2-LG | 5 mg/ml | Acetonitrile | Cayman Chemicals |
|  | AA | 5 mg/ml | Ethanol | Cayman Chemicals |
| Synthetic<br>Cannabinoids | CP55940 | 25 mM | Ethanol | Sigma-Aldrich |
|  | WIN55212-2 | 25 mM | Ethanol | Tocris Bioscience |
|  | AM251 | 25 mM | Ethanol | Tocris Bioscience |
|  | AM630 | 25 mM | Ethanol | Tocris Bioscience |
|  | PF-3845 | 25 mM | Ethanol | Sigma-Aldrich |

For both food preference and food intake assays, flies were placed into feeding chambers, and presented with drug-containing liquid food (5% sucrose, 5% yeast extract) and the respective control food in the glass capillaries, as described below.

### Cannabinoid preference assay

The food preference was measured using the Capillary Feeder (CAFE) assay (Figure 1A), as described in the previous studies ^52,53^. Briefly, eight male flies (3-5 days old) were distributed into each vial containing 1% agarose at the bottom. Four 5 μl glass capillaries (VWR, USA) were inserted into the vial from the lid. Two capillaries were filled with normal control liquid food and the rest with cannabinoid-containing liquid food. The amount of liquid food in the capillaries was measured per day and replaced with fresh food daily for four consecutive days. Two parallel vials void of flies were used as controls in each assay to determine the extent of liquid evaporation from the glass capillaries. The mean amount of evaporation was subtracted from the values obtained for food consumption by the flies. The preference index was calculated as the (consumption of cannabinoid food– consumption of normal control food)/ total consumption.

### Total food consumption

For the assessment of food intake, the behavioral setup was similar to that in the cannabinoid preference assay with the exception that all four glass capillaries (VWR, USA) contained either control food or cannabinoid food in the first 2 days and normal liquid food in the subsequent two days. The total food consumption (μl/fly) was calculated as (food consumption-evaporation loss)/ number of flies.

### Locomotive activity assay

The locomotion of the flies was assessed using an automated climbing assay, as previously described ^54^. Following treatment with cannabinoids for two days, flies were transferred into the climbing vials and were acclimatized for 5 min before testing them. Flies were allowed to climb up the walls of the vials before tapping them down to the bottom at 1-min intervals and this was repeated five times. The testing session was videotaped for analysis and the average climbing height of each fly in the first 6 s of the assay was recorded to determine the locomotive behavior of the flies.

### Starvation resistance assay

Flies pre-treated with cannabinoids for two consecutive days were used to examine starvation resistance. The resistance to starvation was measured as the survival rate of the flies. Ten male flies were placed into each vial containing 1% agarose to prevent desiccation and were monitored at 25 °C and 70% humidity under the 12 h light/ 12 h dark cycle. The number of dead flies was counted every 6-12 h.

### Triacylglycerol Quantification

The levels of triacylglycerol (TAG) were measured in flies according to a previous study ^55^. Following treatment of cannabinoids for two consecutive days, 10 adult male flies per vial were starved for 18 h. These flies were subsequently homogenized in 100 µl of PBS+1% Triton-X and immediately heated at 70 °C for 10 min to inactivate lipases. 20 µl heat-treated homogenates were incubated with the same amount of triglyceride reagent (T2449; Sigma, UAS) or PBST at 37 °C for 60 min. 30 µl of each sample were subsequently added to 100 µl free glycerol reagent (F6428; Sigma, USA) in a clear 96-well plate and incubated for 5 min at 37 °C. The absorbance of the samples was assayed using a multimode microplate reader at 540 nm. TAG concentration was determined by subtracting the absorbance for the free glycerol in the untreated samples from the total glycerol concentration in samples that were incubated with the triglyceride reagent. The levels of TAG were calculated based on the triolein-equivalent standard curve. TAG assay was repeated 3-4 times.

### Statistical Analysis

Statistical analysis was performed with Prism 7.03 (GraphPad Software, USA). Statistical significance was established using Student’s t-test or one-way ANOVA followed by Dunnett’s post-hoc test in the cannabinoid preference and food intake assays, and Log-rank (Mantel-Cox) test in the starvation resistance assays. Differences between treatment groups were considered to be statistically significant at \**p*< 0.05, \*\**p*< 0.01, \*\*\**p*< 0.001. The data were expressed as mean ± standard error of mean (S.E.M).

## Acknowledgments

We thank M. Noll, J.C.P. Yin, and the Bloomington Stock Center (BSC) for generously providing fly stocks. We thank the Yu lab members for helpful discussion.

## Author Contributions

J.H. and F.Y conceived and designed the study. J.H. and A.T. performed most of the experiments. S.N. conducted some of TAG assays. J.H., A.T. and F.Y. analyzed the data. J.H., S.N. and F.Y. wrote the paper.

