## Supplementary Figure 1-3 for "Cannabinoids modulate food preference and consumption in *Drosophila melanogaster*"

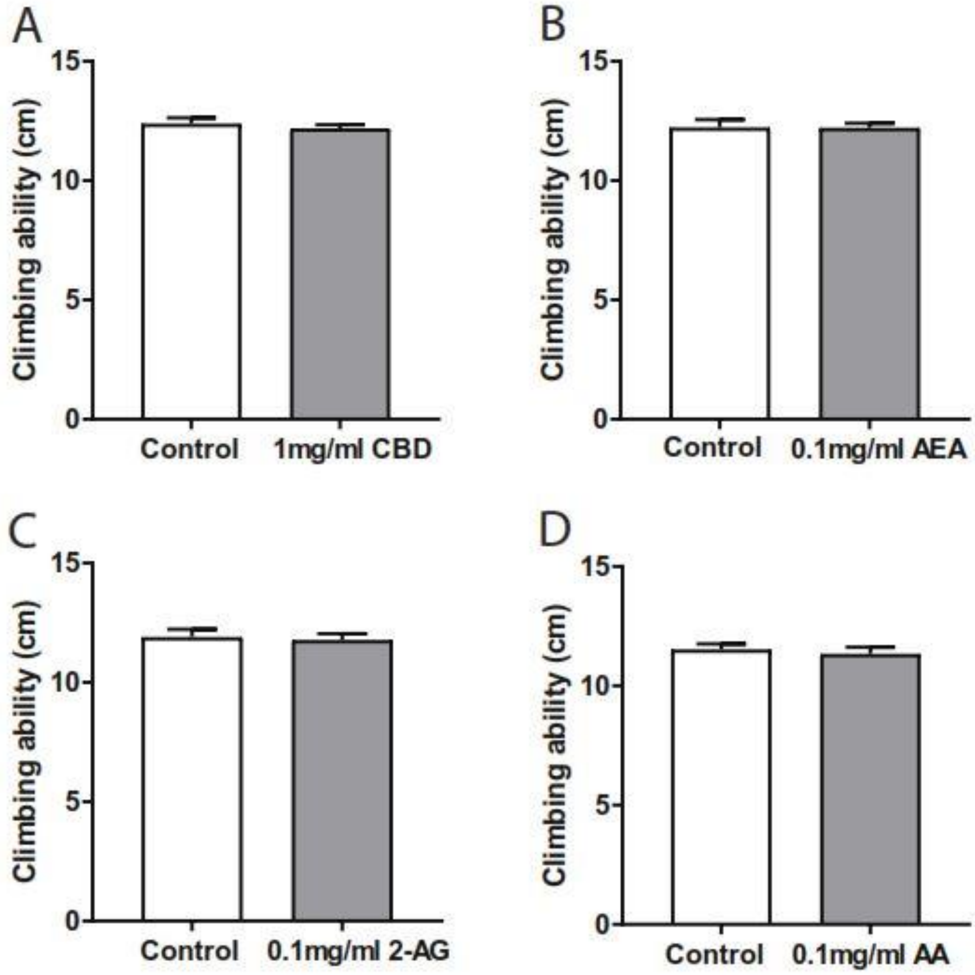


**Figure S1. No or negligible effects of cannabinoids on climbing ability of the treated flies.** Following two days of treatment with cannabinoids, the climbing ability of the flies was assessed to determine the locomotive behavior. Pre-treatment with 1 mg/ml CBD (**A**), 0.1 mg/ml AEA (**B**), 0.1 mg/ml 2-AG (**C**), 0.1 mg/ml AA (**D**) (n=8-12 vials/group) did not alter the climbing performance. Data are represented as mean ± S.E.M.

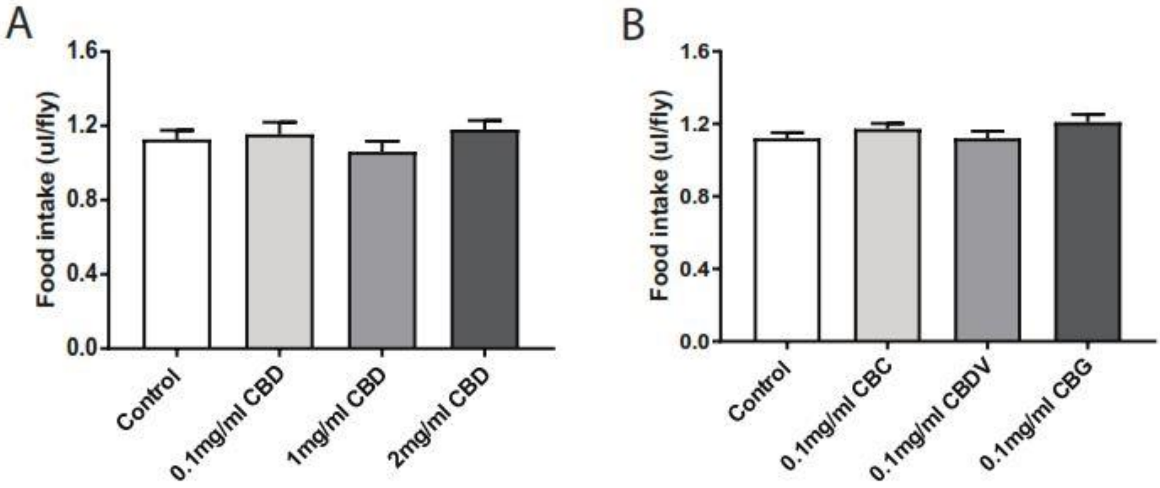


**Figure S2. Initial food intake was not altered during the phytocannabinoids training.** Flies consumed similar amounts of food containing various concentrations of CBD (0.01, 0.1and 1 mg/ml; n=12-19 vials/group) (**A**), or CBDV, CBC or CBG (0.1 mg/ml; n-32-33 vials/group) (**B**) during two days of the training (day -2 and day -1) as compared to the respective control solutions. Data are represented as mean ± S.E.M.

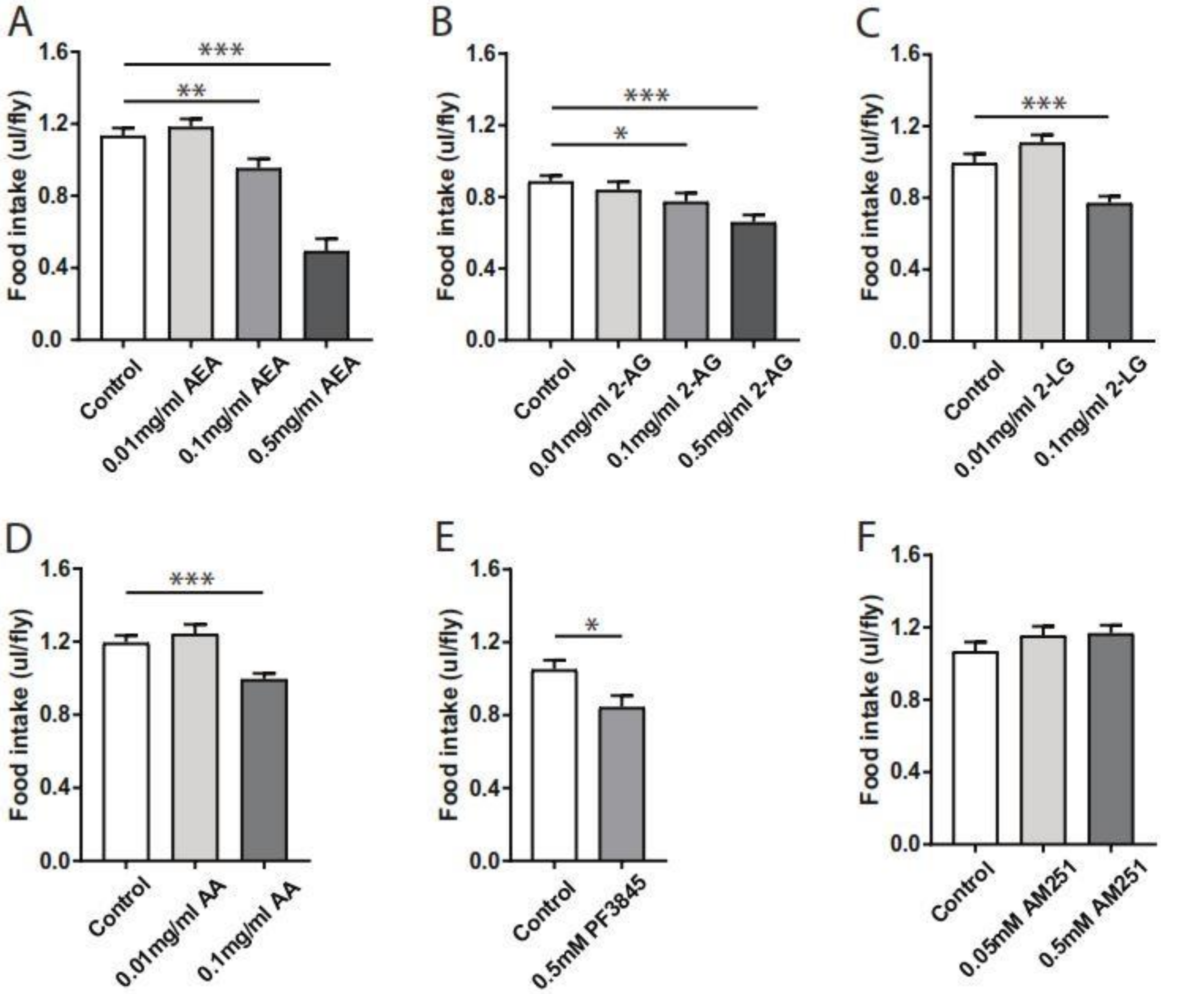


**Figure S3. Endocannabinoids-containing food intake decreased during the initial training.** The amount of food with AEA (0.1 and 0.5 mg/ml) (**A**), 2-AG (0.1 and 0.5 mg/ml) (**B**), 2-LG (0.1 mg/ml) (**C**), AA (0.1 mg/ml) (**D**) and PF3845 (0.5 mM) (**E**) consumed by the flies during two days of the training (day -2 and day -1) was significantly lower when compared to the respective control solutions (n=10-27). The consumption of food with AM251 (**F**) by the flies did not differ from the control groups (n=18 vials/group). Data are represented as mean ± SEM. One-way ANOVA followed by Dunnett's post-test was applied to determine statistical significance. *$p$<0.05, **$p$<0.01, and ***$p$<0.001.
